# Single-cell Transcriptomic Variance Analysis Reveals Intercellular Circadian Desynchrony in the Alzheimer’s Affected Human Brain

**DOI:** 10.64898/2026.03.23.713759

**Authors:** Henry C. Hollis, Anthony Veltri, Ksenija Korac, Vilas Menon, David A. Bennett, Sean M. Ronnekleiv-Kelly, Junhyong Kim, Ron C. Anafi

## Abstract

Bulk tissue rhythms arise from the coordination of thousands of individual cellular oscillations. Bulk rhythm amplitude differences may reflect changes in the amplitude of the underlying cellular oscillators or changes in their temporal coherence. To resolve this fundamental ambiguity, we developed ORPHEUS (**O**scillatory **R**hythm **P**hase **H**eterogeneity **E**stimated **U**sing **S**tatistical-moments), an analytical method that quantifies cellular desynchrony by leveraging the unique 12hr rhythmic signature it imparts on intercellular expression variance. After validating ORPHEUS in silico and on data from the mouse suprachiasmatic nucleus (SCN), we applied it to data from the mouse liver and human brain to uncover disease- and pathway-related differences in intercellular synchrony. In both tissues, we found that circadian synchrony is higher in cells and samples with higher MTORC activity. Most critically, we observed a dramatic loss of cellular synchrony in excitatory neurons from subjects with Alzheimer’s Disease (AD) dementia. By decoupling the influence of cellular amplitude and synchrony, ORPHEUS introduces a new, interpretable tool for analyzing circadian coordination in time-course single-cell data.

## Introduction

Circadian rhythms organize nearly every aspect of our physiology, coordinating processes from sleep-wake cycles to metabolism and immune responses(1). At the tissue level, these rhythms emerge from the coordinated action of millions of individual cellular oscillations, each driven by cell-autonomous molecular clocks(2). In mammals, the suprachiasmatic nucleus (SCN) of the hypothalamus helps synchronize these tissue and cell-type specific cellular clocks(2, 3).

The advent of single-cell RNA sequencing (scRNA-seq) has provided valuable insights into the circadian transcriptome, allowing the characterization of rhythmic gene expression within heterogeneous cell populations(4–8). A standard approach to identify rhythms in these data involves generating “pseudobulk” signals, averaging gene expression of cells of the same type in each sample and timepoint(4–9). However, this approach harbors a critical ambiguity when identifying dampened or low-amplitude rhythms: Such signals could arise from two biologically distinct scenarios: 1) a population of highly synchronized cells that have uniformly dampened amplitudes, or 2) a population of robust cellular oscillators with reduced temporal coherence.

These two scenarios are fundamentally different modes of circadian disruption. For example, a uniform reduction in amplitude may indicate that circadian output genes are receiving a “weaker” signal from the core clock. In contrast, an increase in phase dispersion among cells suggests weaker oscillator coupling, where cell-autonomous rhythms may be functioning normally but the cell-cell communication pathways that synchronize or entrain them are impaired. Moreover, it is possible that different circadian output pathways may gain or lose synchrony as non-circadian driving forces have more or less influence. Disentangling these scenarios is likely crucial for the understanding and treatment of circadian changes associated with aging and disease. Yet, a quantitative framework to decouple the contributions of cellular amplitude and synchrony in time-course scRNA-seq data has been missing.

One approach to quantifying synchrony in these data involves assigning a relative circadian phase to each cell using CYCLOPS(10, 11), Tempo(12), CHIRAL(13), or some other such tool. However, CYCLOPS and CHIRAL were developed for bulk or pseudobulk rhythms, and all these tools often fall short when estimating phases of individual cells given noisy and extremely sparse single-cell transcriptomic data. To fill this gap, we developed ORPHEUS (Oscillatory Rhythm Phase Heterogeneity Estimated Using Statistical-moments). Unlike previous methods, ORPHEUS leverages the analytic relationship between cellular desynchrony, pseudobulk expression, and intercellular variance to provide interpretable, gene-wise estimates of phase dispersion. These gene-wise estimates can then be combined to assess whole cell or pathway specific measures of desynchrony. After validating ORPHEUS in silico and against experimental measurements in the mouse SCN, we apply it to uncover novel, cell-type-specific synchrony patterns in the mouse liver. Finally, we utilize ORPHEUS to characterize the breakdown of circadian coordination in human brains with AD dementia, using the ROSMAP dataset(14).

## Results

### The ORPHEUS Method: Determining Cellular Synchrony from Statistical-Moments

As averages, bulk expression rhythms hide individual cellular nuances. For example, a population of synchronized cells with uniformly low amplitudes can create the same bulk expression rhythm as a population of high-amplitude oscillators that are somewhat out-of-phase with one another. However, while these scenarios produce the same average or bulk expression profile, they produce different patterns in intercellular variance. In a bulk rhythm made from perfectly synchronized cell rhythms, intercellular variance is a constant, independent of circadian time (Fig 1A). In contrast, in a bulk rhythm made from out-of-phase individual oscillators, intercell variance changes with time (Fig 1B). When a circadian transcript is at its peak or trough, its temporal expression profile is nearly flat. Small phase shifts in an individual cell have little effect on expression. At these same times in the circadian cycle, small phase differences *between* cells add little to intercellular variance (Fig 1B, red arrow). However, when transcripts are midway between peak and trough, small circadian phase shifts between cells have a stark effect on expression and intercellular variance increases (Fig 1B, blue arrow). The result is that circadian phase spread produces a 12hr rhythm in intercellular variance, with two peaks and two troughs per 24hr cycle. In transcripts influenced by both non-circadian variation and intercellular circadian phase dispersion, this rhythmic component of variance specifically reflects phase spread and circadian desynchrony (Fig 1C). ORPHEUS leverages the analytical relationship between temporal rhythms in intercellular variance and pseudobulk expression to quantify the spread of individual cell rhythms, or phase dispersion.

**Figure 1:**
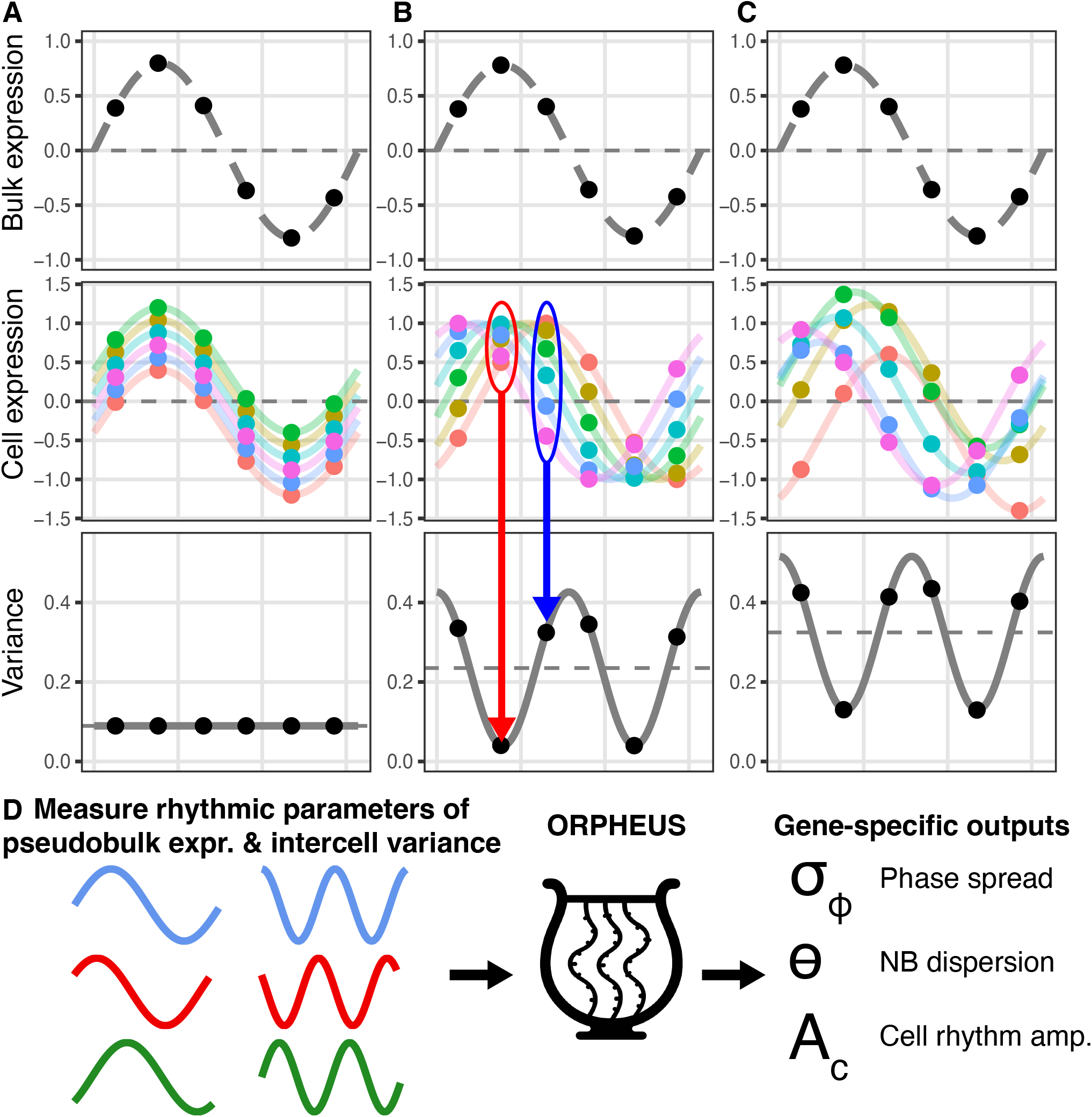
ORPHEUS allows separate estimations of cellular amplitude and synchrony via 12hr harmonic rhythms in intercellular variance. **(A)** Bulk rhythms (top) are an average of individual cell rhythms (middle). When cell rhythms are synchronized, the variance of individual cell expression (bottom) is constant and has no time-dependence. **(B)** Composite bulk rhythm, identical to (A, top), is the result of individual cellular oscillations (middle) with higher amplitude that are somewhat out-of-phase. In this scenario, intercellular variance (bottom) changes with time. Small phase shifts between cells have little effect on intercellular variance at phases near peak or trough expression (red arrow). Near inflection points, phase shifts have a notable effect on variance (blue arrow). The result is a 12hr rhythm in variance (bottom). **(C)** The same bulk rhythm (top) can also arise from individual cellular oscillations (middle) with both non-circadian variation and intercellular circadian phase dispersion. The 12hr rhythmic component of variance (bottom) specifically reflects phase spread. **(D)** ORPHEUS combines the rhythmic parameters from pseudobulk and intercellular variance with the analytic relationship between these signals to solve for gene-specific estimates of phase dispersion, cellular amplitude, and negative binomial overdispersion.

Single-cell transcriptomic data are well described by a negative binomial (NB) distribution as single-cell counts are often overdispersed (mean < variance)(15, 16). We modeled each cell as sharing the same rhythmic parameters for a given transcript (amplitude (*A*_*c*_), MESOR (*G*), and NB overdispersion (*θ*)) but differing in random, normally distributed non-circadian effects 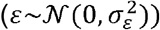 and circadian phase offsets 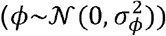. We derived the expected values for the MESOR and amplitude of the pseudobulk rhythms along with the MESOR and amplitudes of the 24hr and 12hr variance rhythms (**Materials and Methods**) as a function of these unknown cellular parameters. ORPHUES estimates the amplitude and phase of expression and variance rhythms directly from observed time-course count data and then solves the system of equations we derived to compute the circadian phase dispersion, cellular amplitude, and NB overdispersion for each circadian transcript (Fig 1D).

### Evidence of Phase Dispersion in Real Single-Cell Data

To validate the core premise of ORPHEUS, we first sought to confirm that 12hr rhythms in intercellular transcriptional variance exist within single-cell time-course data. We re-analyzed public scRNA-seq time-course datasets from mouse SCN(4), mouse liver(17), and human brain(18). Across cell types, species, and datasets, we observed prominent 12hr rhythms in the intercellular variance of transcripts with 24hr expression rhythms, demonstrating that the hypothesized variance signal is real and measurable (Fig 2A-E).

**Figure 2.**
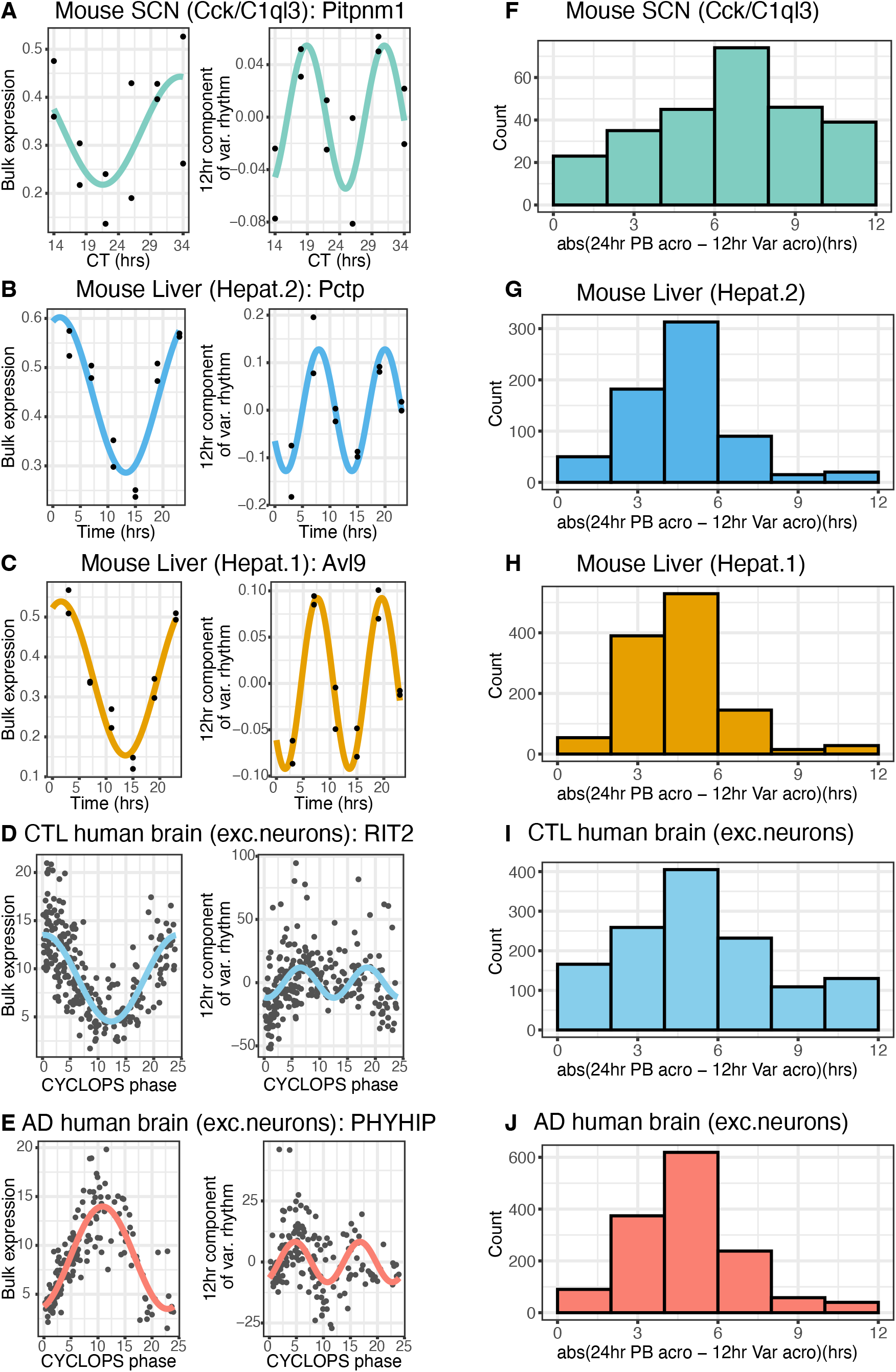
12hr intercellular rhythms are observable across species and tissues. **(A-E)** Illustrative examples of genes with significant 12hr variance rhythms across tissues and species: mouse SCN (*Pitpnm1*), mouse liver (*Pctp, Avl9*), and human DLPFC excitatory neurons (*RIT2, PHYHIP*). In each panel, the 24hr pseudobulk expression (left) is paired with the extracted 12hr component of intercellular variance (right). For mouse SCN **(A)**, data collected over 48 hours is plotted as a function of circadian time (CT) on a 24hr axis. For human data **(D– E)**, individual subjects are ordered by CYCLOPS phase. (**F–J**) ORPHEUS predicts 12hr rhythms in variance arising from phase dispersion will peak 6 hours delayed from the acrophase of pseudobulk expression rhythms. Histograms showing the absolute acrophase difference between the 24hr pseudobulk and 12hr variance rhythms across tissues and species in transcripts with significant rhythms in these signals (**Materials and Methods**).

The theoretical model upon which ORPHEUS is based predicts that 12hr rhythms in variance arise from two distinct sources: 1) intercellular phase dispersion generates 12hr rhythms as explained above. 2) NB overdispersion introduces 12hr rhythms as the variance of an NB-distributed variable, *y*, is a function of both its mean, 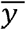, and its square, 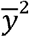. Since the mean expression 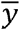 follows a 24hr rhythm, the 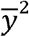 term generates a distinct 12hr component in the variance. Crucially, these two sources generate 12hr rhythms with a predictable phase relationship when compared to bulk expression: 12hr rhythms driven by overdispersion are expected to have peaks aligned with the extremes of the pseudobulk rhythms. Variance due to phase dispersion is minimized at the extremes of the pseudobulk rhythm and thus should be phase-delayed by 6 hours relative to the 24hr pseudobulk expression acrophase. To test this, we calculated the absolute acrophase difference between the 12hr component of intercellular variance and the 24hr pseudobulk rhythms for circadian genes (p<0.05, BooteJTK; BH.q<0.1 & amplitude ratio≥ 0.2, nested cosinor models for human data) with strong evidence of 12hr rhythms in variance (p<0.05 BooteJTK; favorable AIC for human data). We observed a peak at around 6 hours (Fig 2F-J), consistent with the hypothesis that intercellular phase dispersion contributes to the variance dynamics observed in biological signals.

### Validation of ORPHEUS in silico and in Mouse SCN

The accuracy of ORPHEUS’s estimates of phase dispersion depends on both our accuracy in estimating rhythm parameters and the sensitivity of our equations to errors in these measured parameters. We performed simulations to estimate the accuracy of ORPHEUS when ground truth is known. We simulated count data from thousands of cells with both non-rhythmic and rhythmic transcripts, having various cellular amplitudes, phase dispersions, and negative binomial parameters (**Materials and Methods**). In simulations mimicking experiments with high sampling density (24,000 cells across 12 timepoints), ORPHEUS accurately recovered the true phase dispersion across a physiologically relevant range of 1–5 hours (Fig 3A). Reducing the sampling density to a more challenging 4hr interval with fewer cells (8,000 cells across 6 timepoints) increased the spread of gene-wise phase dispersion estimates, but the accuracy of ORPHEUS remained roughly the same (Fig 3B).

**Figure 3:**
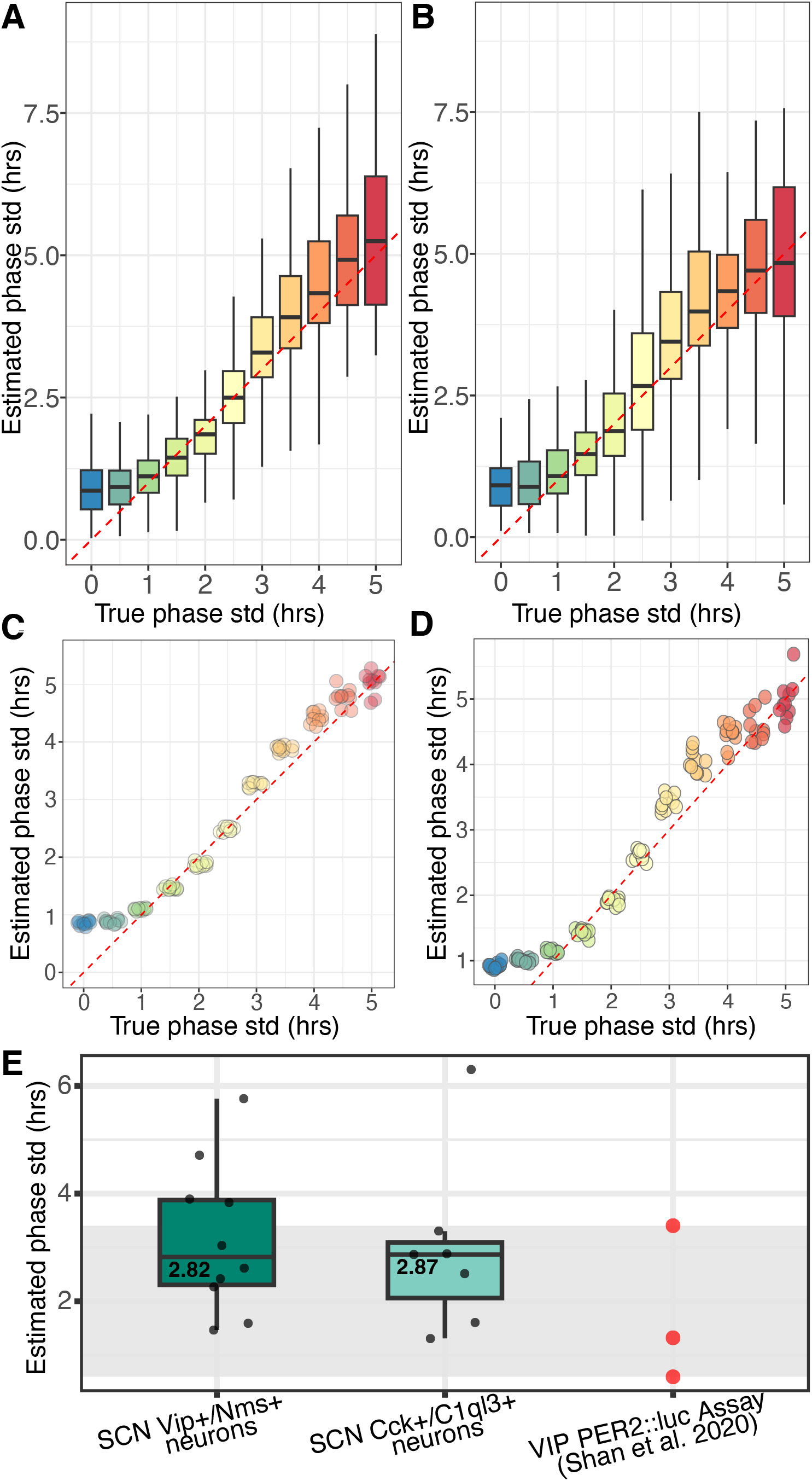
Validation of ORPHEUS in silico and in mouse SCN. We simulated count data from thousands of cells with both non-rhythmic and rhythmic transcripts, having various cellular amplitudes, phase dispersions, and negative binomial parameters. (**A**-**B**) ORPHEUS accurately estimates gene-wise phase dispersion in both highdensity (**A**; 24,000 cells, 12 timepoints) and realistic (**B**; 8,000 cells, 6 timepoints) sampling regimes. **(C-D)** Recovery of “cell-type” (median of gene-wise estimates) phase dispersion parameters from simulated data in both high-density (**C**; 10 independent simulations of 24,000 cells across 12 timepoints) and realistic (**D**; 10 independent simulations of 8,000 cells across 6 timepoints) sampling regimes. The red dashed lines indicate perfect agreement (y=x). (**E**) Biological validation in mouse SCN. ORPHEUS-derived phase dispersion estimates for SCN neurons (green boxplots; derived from time-course scRNA-seq; Wen et al. 2020) are consistent with phase dispersion values measured in SCN slices using *Per2*::LUC imaging (red points; Shan et al. 2020). The gray shaded region highlights the physiological range established by the experimental data. Box plots denote the median (cell-type composite) and interquartile range (IQR) of gene-wise dispersions. Whiskers extend to points within 1.5 × IQR.

We also evaluated the performance of ORPHEUS at the cell-level, using the median of gene-wise estimates as a cell-level measure of phase dispersion. For each true phase dispersion value, we generated 10 independent simulation trials, calculating the cell-level estimate for each trial and comparing it to the ground truth. As expected, the cell-level results are more robust and precise across the tested range of phase dispersions (Fig 3C-D). Cell-level estimates of phase dispersion remained quite accurate with as few as 10 genes.

For biological validation, we turned to measurements of single-cell rhythms in mouse SCN VIP neurons quantified using PER2::LUC bioluminescence imaging(19). We compared the phase dispersion experimentally observed to estimates of phase dispersion derived from the application of ORPHUES to time-course single-cell data from the mouse SCN(4). This dataset features uniformly sampled timepoints, so we utilized the non-parametric BooteJTK algorithm(20) to identify rhythm parameters in pseudobulk and intercellular variance. We supplied these rhythmic parameters to ORPHEUS and estimated cellular phase dispersions of 2.82hrs among VIP+/Nms+ SCN neurons (n=10 gene-wise estimates) and 2.87hrs in Cck+/C1ql3+ SCN neurons (n=7 gene-wise estimates). These de novo estimates are in good accord with the past experimental observations (Fig 3D).

### Assessment of Intercellular Hepatocyte Synchrony in the Mouse Liver

We next estimated intercellular circadian phase dispersion in the mouse liver. As with the SCN data, the uniform temporal sampling of this dataset allowed us to use BooteJTK(20) to estimate gene-wise rhythm parameters. We initially applied ORPHEUS to an aggregated population of all hepatocytes, which revealed a median phase dispersion of 3.09hrs (n=67 gene-wise estimates), comparable to, but slightly higher than, the phase dispersion observed across cell subtypes in the mouse SCN (Fig 4A).

**Figure 4:**
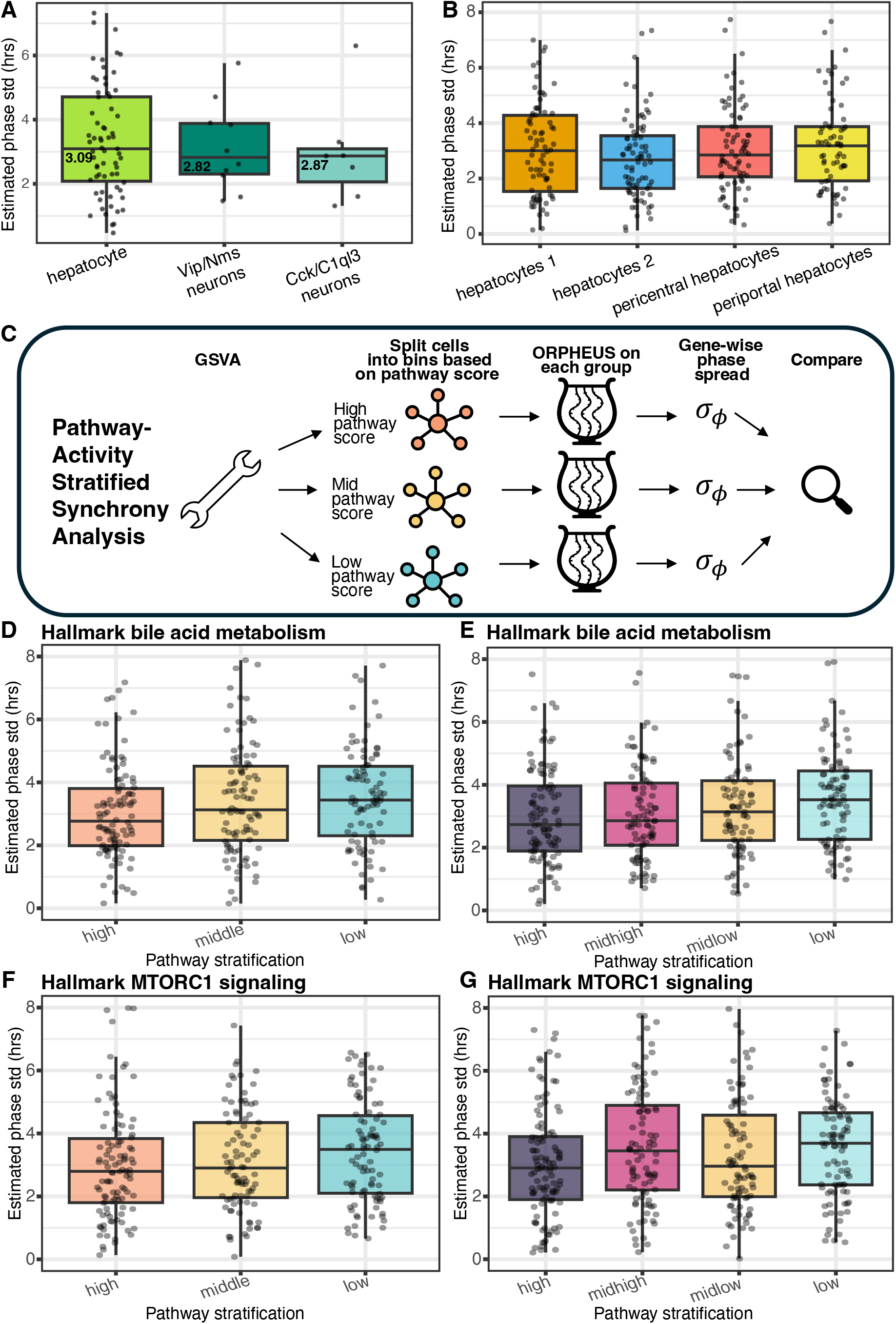
Assessment of hepatocyte synchrony in the mouse liver. (**A**) Gene-wise ORPHEUS estimates of phase dispersion in mouse hepatocytes (Veltri et al., 2025) compared to SCN subtypes. (**B**) Phase dispersion estimates across four annotated hepatocyte subtypes. Synchrony was not significantly different between subtypes as assessed by bootstrapping of cell type labels. (**C**) Schematic of pathway-activity stratified synchrony analysis. Cells are assigned a pathway-activity score using GSVA for each Hallmark pathway. Cells are binned by score (e.g., high, middle, low) within each timepoint and pathway. Phase dispersion is estimated within each bin. Pathways with activity associated with synchrony are identified. (**D**–**G**) **Bile acid metabolism (D, E)** and **MTORC1 signaling** (**F, G**), both predict intercellular synchrony. In both cases, cells with high pathway activity score exhibit significantly higher synchrony (p=0.004; bile acid metabolism, p=0.002; MTORC1 signaling, empirical bootstrapping). This relationship is robust whether cells are stratified into tertiles (**D, F**) or quartiles (**E, G**). Box plots denote the median (cell-type composite) and interquartile range (IQR) of gene-wise dispersions. Whiskers extend to points within 1.5 × IQR.

We then used ORPHEUS to separately assess synchrony in four distinct hepatocyte subtypes as labeled by the original study authors (Fig 4B). Our bootstrap significance procedure (**Materials and Methods**) revealed no statistical difference in phase dispersion between these hepatocyte subtypes (n=68-82 gene-wise estimates).

### Metabolic Drivers of Hepatic Synchrony

The use of ORPHEUS allowed us to probe biological pathways involved in regulating circadian synchrony. At each time-point, we used Gene Set Variation Analysis (GSVA)(21) to score individual cells based on the activity of each Hallmark pathway. For each pathway, we ranked cells by pathway score and stratified them into discrete bins. We then used ORPHUS to quantify synchrony in each bin and identified pathways where synchrony consistently differed across pathway activity bins (Fig 4C).

Hepatocytes with high **bile acid metabolism** scores exhibited significantly lower phase dispersion (higher synchrony) than those with low scores (p=0.004; BH.q=0.088, empirical bootstrapping) (Fig 4D-E). We also observed the same robust relationship for **MTORC1 signaling**: hepatocytes with high MTORC1 activity displayed tighter circadian coordination than their low-activity counterparts (p=0.002; BH.q=0.088, empirical bootstrapping) (Fig 4F-G). The pathway scores for these two processes were only weakly correlated (Pearson r=0.15), indicating that these are distinct biological mechanisms independently associated with circadian precision. These findings remained robust when cells were stratified into either three or four groups, consistently showing that increased pathway activity associated with tighter circadian coupling. As bile acid metabolism shows spatial variation and is highest near the central vein of the portal triad, these results suggest that the liver’s spatial and metabolic gradients may play a role in the regulation of circadian synchrony.

While we didn’t observe a significant difference in synchronization between liver subtypes with the original cell-type labels (Fig 4B), this may be because the cell-type labels were clustered using rhythmic gene expression. Cell-type labels show evidence of circadian corruption as labeled hepatocyte proportion changes over the course of the circadian day despite that these are fairly stable and non-migratory cell types. This suggests that these labels may not reflect robust, biologically distinct cell types.

### Disruption of Circadian Synchrony in Human Brains with Alzheimer’s Disease

One promising use for measuring circadian desynchrony is in understanding circadian dysregulation in human disease. We applied ORPHEUS to a large-scale, single-nucleus RNA-sequencing (snRNA-seq) dataset from the dorsolateral prefrontal cortex (DLPFC) obtained from the ROSMAP consortium(14). We analyzed 409 subjects, including 250 aged individuals without dementia (CTL) and 159 individuals with AD dementia(22). Previous analysis of these data identified a population of excitatory neurons (subtypes 3 & 5) with high circadian signal-to-noise ratio(8). Ordering these data using CYCLOPS 2.0, AD affected subjects demonstrated cell-type specific changes in clock output rhythms. Here, using the circadian phases previously inferred for these DLPFC samples, we applied ORPHEUS to quantify phase dispersion in this neuronal subpopulation with especially robust clock signal (excitatory neurons subtypes 3 & 5). To accommodate the continuous phase assignments and clinical covariates in this human cohort, we employed cosinor regression to estimate the gene-wise rhythm parameters of the pseudobulk and variance time-courses.

### Excitatory Neurons Exhibit Widespread Desynchrony in AD

We observed a significant reduction (p<2.2×10^−16^, Wilcoxen rank-sum test) in circadian synchrony within the AD excitatory neuron population (*σ*_*ϕ*_ =3.76hr, n=526 genes) compared to CTL (*σ*_*ϕ*_ =3.23hr, n=464 genes). This overall loss of coordination was evident in both an unpaired gene-wise synchrony test (Fig 5A) and in direct comparisons of synchrony for the same genes across conditions (Fig 5B; p=5.52×10^−14^, n=278 genes).

**Figure 5.**
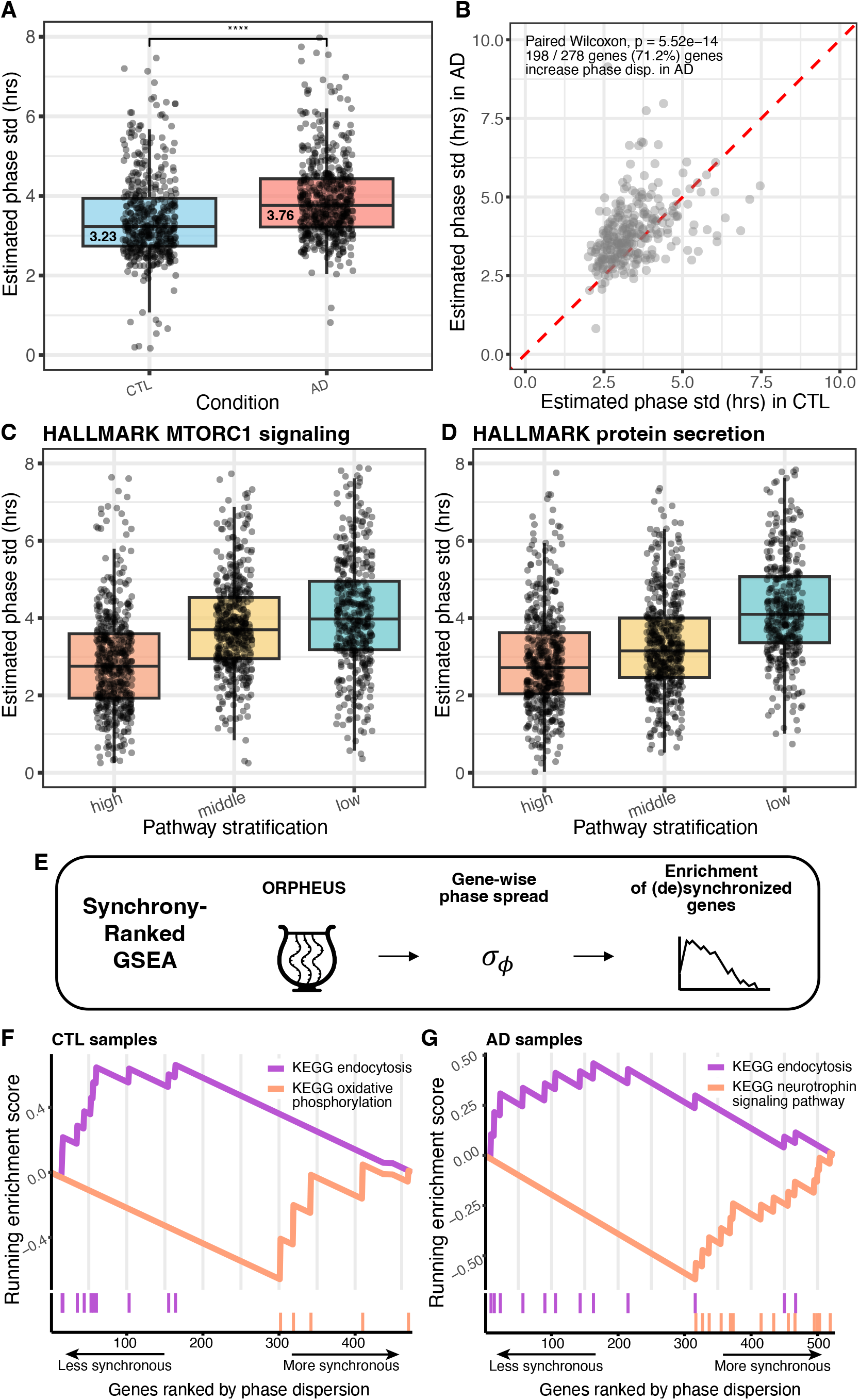
Assessment of inter-neuronal synchrony in human brains with and without AD dementia. (**A**) Gene-wise (points) and cell-type (median) assessment of phase dispersion in excitatory neurons from CTL versus AD subjects. AD neurons exhibit significantly higher phase dispersion (lower synchrony) than controls (*p* < 2.2 *x* 10^−16^, Wilcoxon rank-sum test). (**B**) Comparison of dispersion estimates from transcripts (n=278) that met cycling significance criteria for both expression and variance independently in CTL and AD subjects. Rhythmic transcripts lose synchrony in the AD subjects (*p* = 5.52 × 10^−14^, paired Wilcoxon test; red dashed line = identity). (**C**-**D**) Pathway-activity stratified synchrony analysis within the AD cohort. Subjects with higher activity in (**C**) **MTORC1 signaling** (p=0.032; BH.q=0.26, empirical bootstrapping) and (D) **protein secretion** (p=0.025; BH.q=0.26, empirical bootstrapping) show higher intercellular synchrony. (**E**) Schematic of synchrony-ranked gene set enrichment. Genes are ranked by their estimated phase dispersion values and analyzed via GSEA. (**F**-**G**) Functional enrichment profiles. In both CTL and AD excitatory neurons, **KEGG endocytosis** (purple) is consistently enriched among the most desynchronous circadian transcripts (peaking left). Conversely, **KEGG oxidative phosphorylation** (**F**, CTL) and **KEGG neurotrophin signaling** (**G**, AD) are enriched among the most highly synchronized transcripts (peaking right).

We extended our analysis by stratifying subjects by other metrics of AD severity. Binning all subjects by CERAD score, a semi-quantitative measure of neuritic plaque burden, revealed a significant loss in synchrony associated with increasing plaque density. Moreover, stratification of subjects by Braak stage, which measures the severity and distribution of neurofibrillary tangles, demonstrated a significant loss in synchrony associated with increasing tau pathology.

### Pathway-Activity Stratified Synchrony Approach Reveals Association Between MTORC Pathway and Circadian Synchrony

Unlike the mouse liver data where we stratified *cells* by pathway activity score, here we scored and stratified *subjects/samples* by GSVA, as we are interested in subject-wise differences in pathway activity. We did this analysis independently for CTL and AD subjects. As before, we performed ORPHEUS independently on each pathway bin and compared phase dispersion metrics.

Among CTL subjects, there were no pathways for which stratifying subjects by pathway score yielded significant differences in synchronization. In AD, stratifying subjects by either **MTORC1 signaling** (p=0.032; BH.q=0.26, empirical bootstrapping) or **protein secretion** (p=0.025; BH.q=0.26, empirical bootstrapping) activity yielded significant differences in estimated synchronization (Fig 5C-D). The relationship between pathway activity and synchrony persisted if subjects were stratified into 2 or 3 bins. Strikingly, the identification of MTORC1 signaling as a positive regulator of synchrony in human neurons mirrors our finding in mouse hepatocytes, pointing to a conserved correlation between this anabolic hub and circadian coherence across distinct tissues and species. The pathway scores for MTORC1 and protein secretion were moderately correlated (Pearson r = 0.49).

With the ample number of cells and subjects in this dataset, we were able to identify many rhythms in pseudobulk and variance time-courses and subsequently estimate the phase dispersion for many genes via ORPHEUS. This allowed us to perform synchrony-ranked gene set enrichment analysis (Fig 5E): Genes were ranked based on their ORPHEUS-derived phase dispersion estimates and then input to Gene Set Enrichment Analysis (GSEA)(23, 24) to identify pathways enriched among relatively synchronous and desynchronous genes. We found in CTL samples that KEGG oxidative phosphorylation (p=0.018, BH.q=0.201) and Hallmark glycolysis (p=0.038, BH.q=0.295) were enriched among the most synchronous genes. Endocytosis was significantly enriched among the relatively desynchronous genes (p<0.001, BH.q=0.012) (Fig 5F). In AD samples, KEGG neurotrophin signaling was significantly enriched among the most synchronous genes (p<0.001, BH.q<0.001). Directionally consistent to CTL samples, KEGG endocytosis showed a trend for enrichment among the more desynchronous genes (p=0.037, BH.q=0.56) (Fig 5G).

Previously, we found that in the brains of AD affected subjects, genes in the oxidative phosphorylation (OXPHOS) and ribosome pathways exhibited dampened rhythm amplitudes as compared to CTL(8). A central question remained: Did disease status influence output pathway amplitude in individual cells (uniform dampening) or did these output rhythms lose synchrony? While the stringent filtering for significant fits for both expression and variance rhythms limits the number of OXPHOS and ribosome genes available for direct phase dispersion estimates, the subset of genes that could be modeled provide a mechanistic clue. As a group, circadian phase dispersion for OXPHOS transcripts was higher in AD subjects (*σ*_*ϕ*_ =3.17hr, n=10 genes) compared to CTL (*σ*_*ϕ*_ =2.76hr, n=5 genes) (p= 0.005, Wilcoxen rank-sum test). A difference in phase dispersion between conditions for ribosome genes is also apparent, though not statistically significant. The cellular amplitude of both OXPHOS and ribosome transcripts was comparable between conditions. These data support the hypothesis that the dampening we observed in bulk rhythms of OXPHOS and ribosome component genes may be partially driven by a loss of intercellular coherence.

## Discussion

In this study, we address a long-standing challenge in circadian biology: the inability to distinguish between changes in cellular amplitude and cellular desynchrony. We derived the governing equations relating circadian dispersion, intercellular variance, and transcript expression rhythms. We demonstrate that 12hr variance rhythms are not only theoretically expected but experimentally observable in time-course single-cell data. To operationalize this, we developed ORPHEUS as an open-source R package that estimates phase dispersion, cellular amplitude, and negative binomial overdispersion for rhythmic genes. By exploiting the relationship between the first and second statistical moments of time-course gene expression (rhythms in mean and variance), ORPHEUS can estimate phase dispersion without having to assign phases to individual cells. Following in silico validation and experimental benchmarking in the mouse SCN, we applied ORPHEUS to data from the mouse liver and the human brain with and without Alzheimer’s Disease. We discovered that excitatory neurons in AD exhibit significantly lower synchrony than those in cognitively normal controls. Crucially, this suggests that the dampening of oxidative phosphorylation and ribosomal rhythms previously observed in human AD subjects can be attributed, at least in part, to elevated phase dispersion rather than a pure loss of cell-autonomous oscillator amplitude.

Stratifying cells and samples by pathway activity, our analysis suggests that synchronization appears to be associated with metabolic state, and in the liver may also reflect anatomic zonation. In both mouse hepatocytes and human excitatory neurons, we observed that MTORC1 signaling, a master driver of protein synthesis, was significantly associated with high circadian synchrony. This computational observation aligns with existing experimental literature identifying the mTOR pathway as a regulator of clock robustness. Previous studies have demonstrated that mTOR signaling is essential for photic entrainment and the maintenance of intercellular synchrony within the SCN; specifically, targeted ablation of mTOR in VIP neurons weakens clock synchrony(25). Furthermore, in peripheral oscillators like the liver, pharmacological and genetic activation of mTOR augments bulk tissue clock amplitude(26). The conservation of this relationship across distinct species and tissues suggests a fundamental link between metabolic activity and clock coherence. High anabolic output may impose a constraint on cell synchronization: energetically expensive processes like protein synthesis or bile acid production may require tight temporal coordination to maximize efficiency(27, 28).

These observations provide support for hypotheses generated in our prior work. Previously, we adapted a mathematical model of the circadian clock to simulate the effects of neurodegeneration on oscillator fidelity. Introducing translational noise and slowing, designed to mimic lowered ribosomal protein abundance observed in AD brains(8, 29–31), resulted in decreased clock precision. Our ORPHEUS analysis found that AD subjects exhibiting lower MTORC1 and protein secretion activity (master regulators of mRNA translation and proteostasis) exhibit reduced intercellular circadian synchrony. While these associations are not necessarily causative, the convergence of our prior modeling with these new observations supports the hypothesis that translational deficits may be important in AD circadian disruption.

Our synchrony-ranked gene set enrichment analysis of human neurons allowed us to analyze the relative synchrony of different clock output pathways and revealed that endocytosis is enriched among the most desynchronous rhythmic transcripts. Endocytosis, important for synaptic vesicle recycling, is likely driven by local, stochastic neuronal firing events. While these endocytosis genes exhibit a population-level rhythm (likely set by both the clock and the sleep-wake cycle), their moment-to-moment expression is tied to heterogeneous firing states. Thus the conclusion that the circadian genes in this clock controlled pathway exhibit high cell-to-cell phase variability is biologically plausible. This finding also highlights the idea that circadian synchrony may vary across biological pathways, reflecting differences in the extent to which noncircadian regulatory processes influence gene expression.

While we believe our application of the concepts underlying ORPHEUS is novel within chronobiology, the coupling of rhythmic expected values and periodic variances is an established analytical principle in other disciplines. In econometrics, for example, models of periodic heteroscedasticity have been employed to evaluate seasonal volatility, capturing how the variance of an asset’s price cycles in tandem with the rhythms of its mean value(32).

Similarly, in climatology, the rhythmicity of a periodic mean and its variance is utilized to model weather predictability(33).

## Strengths and Limitations of the ORPHEUS Framework

A primary strength of our approach lies in its interpretability. Unlike many “black box” machine learning methods, ORPHEUS operates by solving a transparent system of equations derived from a fundamental model of cellular rhythms. The underlying expression and variance rhythms can be directly visualized, and researchers retain full control over the statistical cutoffs used to filter genes. Moreover, ORPHEUS is highly flexible; it allows users to supply cycling results from their preferred algorithms. For instance, using cosinor regression allows researchers to accommodate complex experimental designs and control for technical covariates before phase dispersion is estimated.

Using pseudobulk and variance from hundreds or thousands of cells at each timepoint, ORPHEUS is less affected by the inherent noise and dropout rates common in single-cell expression data. This provides an advantage over single-cell phase-assignment tools (e.g., TEMPO), which can struggle when core clock genes exhibit high dropout rates. For example, in the mouse SCN dataset we re-analyzed, 93% of neurons, 96% of astrocytes and 98% of microglia had 0 reads associated with core clock transcripts (median percentage across core clock genes in each cell type). Likely as a result, the use of TEMPO to assign phases to individual cells and thus quantify phase dispersion results in estimates of desynchrony that vary in time and exceed experimental observation(19). Of course, as sequencing techniques improve, these methods may have increasing utility. ORPHEUS also benefits from its gene-wise estimation of phase dispersion, which enables the identification of distinct levels of circadian coordination within different pathways in the same cellular population.

Finally, the quantitative alignment between ORPHEUS estimates and existing experimental evidence of phase dispersion in the mouse SCN serves as a powerful proof-of-concept for its accuracy.

While ORPHEUS provides a novel framework for quantifying cellular synchrony, our approach incorporates several assumptions and methodological choices. These assumptions may limit the applications of ORPHEUS but also represent important areas for future development. The algorithm currently relies on single-cell time-course data with relatively fine sampling to detect and accurately describe 12hr rhythms in intercellular variance. Various benchmarking studies in circadian rhythm detection have highlighted the concept that multiple samples are required per cycle and more frequent sampling is needed to accurately detect 12hr rhythms as compared to 24(34, 35). While such finely sampled datasets are currently limited, we anticipate a significant increase in their availability as single-cell sequencing becomes more cost-effective.

The mathematical model underlying ORPHEUS incorporates some simplifying assumptions, most notably that the amplitude of rhythms is functionally constant across cells. This assumption is justifiable given that amplitude and period of cellular oscillators are properties defined by the biochemical kinetics of the core clock transcription-translation feedback loop. Because these kinetics are governed by the physical properties of the molecules themselves, they are expected to be largely conserved across a homogeneous cell population. From a dynamical systems perspective, variations in initial conditions or transient biochemical noise only alter the oscillator’s position along the limit cycle (phase) and not the period or amplitude of the oscillator(36–38). Relaxing this assumption may be possible but would create an underdetermined system (fewer equations than variables), requiring the development of a composite optimization strategy (e.g., fixing phase dispersion across genes) for a mathematically tractable problem.

Our in silico simulations show that ORPHEUS is accurate within a specific window of phase standard deviations (1-5 hours). This is to be expected as extreme synchrony eliminates the 12hr variance rhythms and extreme desynchrony flattens both pseudobulk expression and variance rhythms and is poorly modeled by a normal distribution.

As pseudobulk expression rhythms are not perfectly sinusoidal, higher-order harmonics in the expression signal could also generate 12hr variance rhythms. To mitigate this concern and its impact on ORPHEUS phase dispersion estimates, our preprocessing pipeline regresses out the contribution of the full pseudobulk signal in the variance signal.

Currently, comparing phase dispersion between biological groups relies on calculating gene-wise phase dispersions within each group and then testing for differences in central tendency. A more statistically robust future approach may involve a unified model that fixes phase dispersion across genes and formally tests whether adding a “condition/group” variable significantly improves the overall model fit. Such an approach is well established in differential rhythm analysis(39).

A notable limitation in our application of ORPHEUS to the mouse liver dataset is the lack of true biological replicates at each timepoint, a common constraint in time-course scRNA-seq experiments due to high experimental costs. To mitigate this and ensure robust rhythm detection, we adopted a pseudo-replication scheme inspired by the original authors(17), and recent benchmarking simulations(9). While we tried to make this as robust as possible (**Materials and Methods**), this scheme cannot fully substitute for a well sampled experiment with biological replicates. Consequently, we utilize these liver data primarily to demonstrate the principles and analytical potential of the ORPHEUS framework. The biological relationships identified here serve as hypothesis-generating observations that warrant future validation in properly replicated multi-subject cohorts.

While ORPHEUS is agnostic to upstream rhythm-detection methods, benchmarking to determine which upstream algorithm yields the most accurate inputs given different experimental designs would likely be of value. In this study we use BooteJTK to identify rhythms in datasets with uniform collection times, as it has been shown to possess high sensitivity for detecting rhythms in sparse data(20) and allows the user to specify a search for rhythms with particular acrophases, unlike the original JTK_CYCLE algorithm(33). To accommodate the continuous phase assignments and clinical covariates in the human brain data, we employed cosinor regression and nested models to identify rhythms in pseudobulk and intercellular variance.

Comparison of ORPHEUS estimates with experimentally observed synchrony in the mouse SCN provided a useful baseline. However, additional validation against benchmarks in which phase dispersion is directly manipulated will be required to fully characterize the algorithm’s limits.

The temporal analysis of transcriptional variation and the ORPHEUS package provides a new analytical lens for chronobiology. By decoupling cellular amplitude from intercellular coordination, this framework elevates synchrony to a measurable, experimentally interpretable variable. As time-course single-cell transcriptomics become increasingly accessible, tools like ORPHEUS will be essential for uncovering the mechanistic drivers of circadian breakdown, paving the way for targeted chronotherapeutic interventions in aging and neurodegeneration.

## Materials and Methods

### ORPHEUS

We developed ORPHEUS (Oscillatory Rhythm Phase Heterogeneity Estimated Using Statistical-moments) to quantify cellular synchrony from time-course single-cell transcriptomic data. We model the counts of a gene in a single cell, *S*(*t*), as a random draw from a negative binomial (NB) distribution:

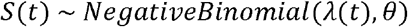

The mean of the distribution, *λ*.(*t*), is a stochastic rhythm defined as:

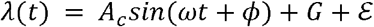

Here, *A*_*c*_ is the cellular amplitude, *ω* is the frequency (fixed at 2*π*/24), G is the MESOR (Midline Estimating Statistic of Rhythm), and *θ* is the NB dispersion parameter, which quantifies intrinsic transcriptional noise. The terms *ϕ* and *ε* represent sources of intercellular heterogeneity: *ϕ* is a random phase offset, assumed to be normally distributed 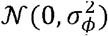, and *ε* is additive baseline noise, distributed 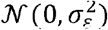. Our goal is to estimate the key unknown parameters, particularly the phase dispersion *σ*_*ϕ*_, cellular amplitude *A*_*c*_, and intrinsic dispersion *θ*.

### Derivation of the Mean and Variance of Rhythmic Gene Counts

To connect our model to measurable data, we derived the expected mean (*ε*[*S*(*t*)]) and variance (*Var*[*S*(*t*)]) for a population of cells at time t. Using the law of total variance,

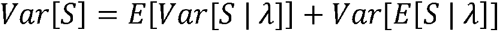

and the properties of the NB distribution,

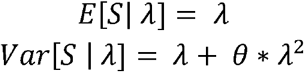

we arrive at the master equation:

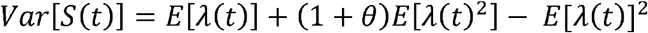

To get an equation for *Var*[*S*] in terms of the variables of interest, we need to derive *E*[*λ* (*t*)] *and E*[*λ* (*t*)^2^]:

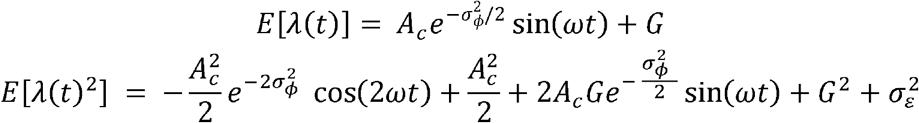

Substituting these into the master equation yields the full expression for *Var*[*S*], which is a superposition of a non-oscillatory DC component, a fundamental 1ω (24hr) component, and a second-harmonic 2ω (12hr) component:

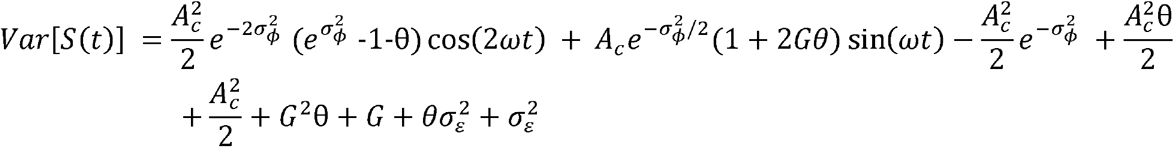

### Estimation of Rhythmic Parameters from Time-Course Data

We estimated the rhythmic parameters of the mean and variance signals directly from processed expression data. For a given gene, we first calculated the mean and variance time-series. For datasets with non-uniform sampling and clinical covariates (human brain data), we employed cosinor regression and nested linear models to identify significant rhythms in these signals. For datasets with uniform sampling, we utilized the BooteJTK algorithm with a custom pipeline to identify 24hr and 12hr rhythms in pseudobulk expression and intercellular variance.

### Rhythm Detection via BooteJTK

For data with uniform sampling, we identified rhythms in both pseudobulk expression and intercellular variance using BooteJTK. BooteJTK has been shown to possess high sensitivity for detecting rhythms in sparse data(20) and allows the user to specify a search for rhythms with particular acrophases, unlike the original JTK_CYCLE algorithm(40). For pseudobulk signals, we specified a search for rhythms with 24hr periods using symmetric reference waveforms, testing for acrophases in the range of 0-23 hours at 1hr intervals.

The detection of 12hr rhythms in intercell variance is more challenging because the signal contains a dominant 24hr component. While these frequencies are theoretically orthogonal, the rank-based nature of the BooteJTK algorithm means that a high-amplitude 24hr oscillation can obfuscate the 12hr harmonic. To isolate the 12hr signal in variance, we subtracted the 24hr and non-oscillatory components of variance from the total variance signal, and provided the resulting “residual variance” signal to BooteJTK.

Specifically, the variance signal *V*(*t*) was modeled as:

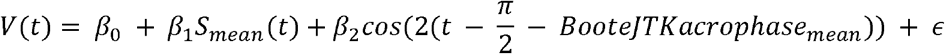

Where *β*_*θ*_ represents the non-oscillatory intercept, *S*_*mean*_(*t*) is the pseudobulk signal (24hr rhythm component), and 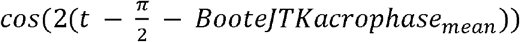 fits the 12hr harmonic (using the acrophase of the mean rhythm identified by BooteJTK, *BooteJTKacrophase*_*mean*_). We then derived a “residual variance” signal by subtracting the fitted non-oscillatory and 24hr components from the total intercellular variance signal:

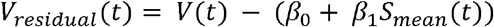

To identify 12hr rhythms in this residual variance signal we performed a targeted BooteJTK search, restricted to genes with significant pseudobulk rhythms (p<0.05, BooteJTK). The equations underlying ORPHEUS hypothesize that 12hr variance rhythms have acrophases either equal to the pseudobulk phase or delayed by 6hr. To accommodate our coarse sampling resolution (4hr), we expanded this search to a window of ±2hr around the predicted phases. Because this approach necessitated a gene-specific list of acrophases to test, BooteJTK was run in basic mode (-B) for each gene individually, searching for rhythms with 12hr periods using symmetric reference waveforms.

Genes identified as rhythmic in both pseudobulk expression (p<0.05, BooteJTK) and variance (p<0.05, BooteJTK) were selected for final parameter estimation. To obtain accurate amplitude and phase estimates, we performed cosinor regression on these genes using functions within the ORPHEUS package. This step was necessary as BooteJTK reports amplitude as the simple difference between the maximum and minimum values rather than a fitted waveform amplitude.

### Rhythm Detection via Cosinor Regression and Nested Models

For data with non-uniform sampling and known clinical covariates (human brain data), we identified rhythms in both pseudobulk expression and intercell variance using cosinor regression and nested models.

### Identifying Pseudobulk/Mean Rhythm Parameters (*DC*_*mean*_, *Amp* _*mean*_, *acrophase*_*mean*_)

For each gene, we fit the pseudobulk/mean signal, *S*_*mean*_ (*t*), to two nested models and used the F test to determine the if the mean signal is significantly rhythmic (BH.q<0.1 and amplitude/MESOR ≥0.2). The F test here has the null hypothesis that *β*_4_=*β*_5_=0.

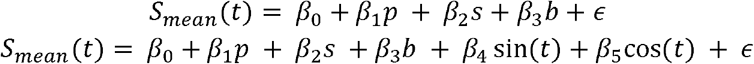

Where *p, s, b* control for subject post-mortem interval, sex, and sequencing batch, respectively. We can extract 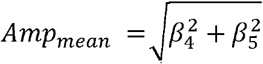, and *acrophase*_*mean*_ = *atan*2 (*β*_4_=*β*_5_) *mod* 2*π*, from the full model. The baseline expression level, or *DC*_*mean*_, was calculated as the mean of the non-oscillatory components of the full model:

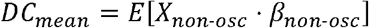

Where *X*_*non-*-*osc*_ represents the design matrix of non-oscillatory variables (intercept and covariates) and *β* _*non-*-*osc*_ represents their corresponding fitted coefficients.

### Identifying 24hr and 12hr Variance Rhythms (*AmpFundamental* _*var*_, *Amp2Harm* _*var*_)

Like the mean signal, we fit the variance signal of a gene, V(t), to two nested models to determine if the variance has significant 12hr rhythms:

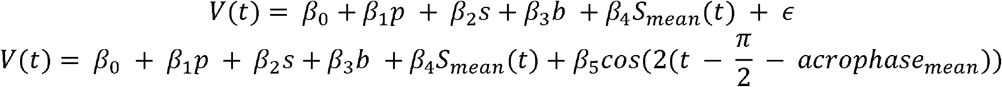

Where *S*_*mean*_ (*t*) still indicates the mean/pseudobulk signal at time t. The full model tests for 12hr rhythms at specific acrophases. Our derived equations predict 12hr rhythms arise in V(t) from both the negative binomial overdispersion parameter (*θ*), and phase dispersion (*ϕ*). The NB overdispersion generates 12hr rhythms in-phase with the 24hr mean rhythm. Phase dispersion generates 12hr rhythms 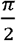 radians (6hrs) offset from the mean rhythms’ acrophases. The amplitude of the 12hr rhythm that we measure in V(t)is the sum of these two 12hr rhythms. *Amp*2*Harm* _*var*_ is the signed amplitude of the 12hr component of the variance timeseries, and is determined by the coefficient of the sinusoid in the regression model above: *Amp*2*Harm* _*var*_= *β*_5_.

Unlike our test for 12hr rhythms with BooteJTK, which has been shown to possess higher sensitivity for detecting rhythms in sparse data compared to cosinor approaches(20), our use of cosinor regression is inherently more conservative and identifies fewer 12hr variance rhythms. To address this, we employed an information-theoretic approach for model selection rather than a strict F-test p-value threshold. For each gene, we calculated the Akaike Information Criterion (AIC) for both the full variance model (which includes the 12hr harmonic term) and the partial null model. A gene was considered to exhibit evidence of a 12hr variance rhythm, and was thus retained for phase dispersion estimation, if the inclusion of the harmonic term improved the model fit sufficiently to yield a lower AIC (*AIC*_.*full*_<*AIC*_*partial*_).

Quantifying the amplitude of the 24hr rhythmic component of variance, *AmpFundamental* _*var*_, requires a secondary step. We fit the “mean-driven” component of variance from the full variance model above, *β*_4_*S*_*mean*_ (*t*), to a linear model (locked to the previously identified pseudobulk acrophase, *acrophase*_*mean*_):

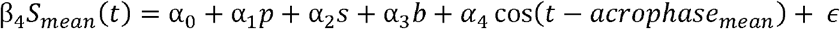

Here, *p, s*, and *b* represent the post-mortem interval, sex, and sequencing batch covariates, respectively. The signed amplitude of the 24hr variance oscillation, purified of technical confounders is determined by the coefficient of the sinusoid in the regression model above: *AmpFundamental* _*var*_=*α*_4._

### Solving for Model Parameters via Method of Moments

By equating the theoretically derived coefficients of the Var[S(t)] and E[S(t)] waveforms with the empirically measured parameters, we obtain a solvable system of equations:

- *G = DC*_*mean*_
- *θ* = ((*AmpFundamental_var_/Amp_mean_*) −1)/2*G*
- Let R = *Amp*2*Harm* _*var*_/*Amp*_*mean*_^2^ then the phase dispersion is: 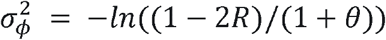
- The cellular amplitude, Ac, is: 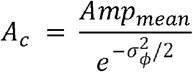

### Single-Cell RNA-Seq Datasets

#### Mouse Liver Data

We re-analyzed single-cell, time-course data from mouse liver(17). While the full dataset consists of 34921 cells, our analysis focused on the hepatocyte population (n=32515). Cell-type clustering and annotations were retained from the original publication. Samples were collected over six discrete timepoints, spaced 4 hours apart over a 24hr period. Initially, all hepatocytes were aggregated into a single population to identify high-confidence rhythmic genes. Because the data were uniformly sampled, we utilized BooteJTK to detect oscillations in both pseudobulk expression and intercell variance. We employed a pseudoreplication strategy where at each timepoint, hepatocytes were randomly partitioned into two independent pseudo-replicates. This partitioning and the subsequent rhythm-detection process was repeated four independent times (4 folds). Genes exhibiting significant rhythms (p<0.05 BooteJTK, for both 24hr pseudobulk and 12hr variance rhythms) across all four folds were retained. These high-confidence rhythmic genes served as input for ORPHEUS to estimate global hepatocyte phase dispersion. In our subsequent analysis of distinct hepatocyte subtypes, we used the same gene set as input to ORPHEUS. While rhythmic genes were identified in pseudoreplicated data, the ORPHEUS estimates for phase dispersion were calculated using the full, non-pseudoreplicated data.

#### Mouse SCN Data

We re-analyzed a previously published single-cell RNA-sequencing dataset of the mouse suprachiasmatic nucleus(4). We analyzed 10,572 neurons with author-provided SCN subtype annotations, collected over a 48hr time-course with a sampling interval of 4hr. The experimental design involved three distinct sampling batches, where samples separated by 12hr were processed in the same batch. Consequently, the batch assignment itself exhibits a 12hr periodicity. This design creates a perfect collinearity between the batch effect and any potential biological 12hr rhythm, rendering it impossible to include a batch term in cosinor regression models without absorbing the 12hr signal of interest. To mitigate this technical confounder while preserving biological variation, we applied ComBat(41) batch correction to the normalized count matrix. Following correction, we aggregated the data to generate pseudobulk expression and intercellular variance time-courses for each gene and performed rhythm detection with BooteJTK.

#### Simulated Data

To validate ORPHEUS, we simulated single-cell RNA-sequencing count data using a negative binomial model incorporating intercellular desynchrony. The expression count y_*gc*_ for gene _*g*_ in cell was drawn from a negative binomial distribution:

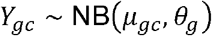

where *θ*_*g*_ represents the gene-specific overdispersion parameter. The mean expression, *μ*_*gc*_, for rhythmic genes was modeled as a cosine function subject to phase noise:

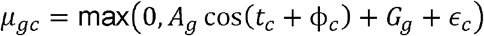

Here, *t*_*c*_ represents the collection time, *A* _*g*_ is the cellular amplitude of gene g, and *G*_*g*_ is the cellular MESOR (baseline expression) of gene g. To simulate phase spread, a cell-specific phase shift ϕ _*c*_ was drawn from a Gaussian distribution, 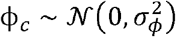, where *σ*_*ϕ*_ represents the “ground truth” phase dispersion parameter to be estimated. Additionally, a modest level of cell-specific Gaussian noise (*ϵ*_*c* ~ 𝒩 (_0,0.3^2^)) was added to the mean expression to mimic variability beyond biological overdispersion.

We evaluated the performance of our algorithm across a range of true phase standard deviations (*σ*_*ϕ*_) under two distinct experimental conditions (sparse and dense sampling regimes):

1. **Mouse Liver Proxy:** To approximate the characteristics of our biological dataset, we simulated *n* = 8,000 cells and *m* = 10,000 genes sampled across 6 time points (4hr resolution). Parameters were drawn from distributions resembling empirical liver data (hepatocytes 1 cell type): amplitude ratios (*A*/*G*) between 0.15-0.75, MESOR (*G*) values between 0.1-5, and dispersion (*θ*) between 0.05-1.5. The proportion of rhythmic genes (*p*_*circ*_) was set to 0.1.
2. **Idealized Experiment:** To test the theoretical limits of the method under high-density sampling, we simulated an idealized dataset with *n* = 24,000 cells sampled across 12 time points (2hr resolution). Gene parameters (*A*/*G, G,θ*) and rhythmic proportions were identical to the mouse liver proxy.

### Human Alzheimer’s Disease Data

We re-analyzed single-nucleus RNA-sequencing data from the dorsolateral prefrontal cortex of participants in the ROSMAP cohort(14), as previously curated and described in our prior work(8). All ROSMAP participants enrolled without known dementia and agreed to annual detailed clinical evaluation and brain donation at death. Both studies were approved by an Institutional Review Board of Rush University Medical Center. Each participant signed informed and repository consents and an Anatomic Gift Act. Our analysis included 409 subjects: 250 aged controls and 159 individuals with AD dementia. Diagnostic labels for our primary analysis (cogdx) were based on clinical diagnosis proximate to death: no dementia (CTL) and Alzheimer’s dementia (AD)(22, 42, 43). We also analyzed samples stratifying by CERAD score and Braak stage(44). Cell-type annotations were retained from the original publication introducing these data(18). To reconstruct temporal information from this static population data, we informatically ordered subjects using the CYCLOPS 2.0 algorithm(10, 11).

## Data Preprocessing and Moment Calculation

To account for technical differences in sequencing depth while preserving the negative binomial structure of single-cell data, we employed a modified Trimmed Mean of M-values (TMM) normalization strategy. First, we filtered out lowly expressed genes, retaining only those with at least 1 count in 1% of cells (or 0.1% for the sparser mouse SCN dataset). Normalization factors were calculated using the calcNormFactors function in the edgeR package(45). Rather than scaling counts to a fixed “per million” constant (CPM), which can arbitrarily shift the magnitude of the data, we scaled counts to the mean effective library size of the dataset. For each sample *j*, an effective library size (*S_j_*) was calculated as the product of the raw library size and the TMM normalization factor. We then derived a sample-specific scaling factor:

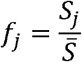

Where 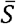 represents the mean effective library size across all samples in the experiment. The normalized count for gene in sample was then defined as:

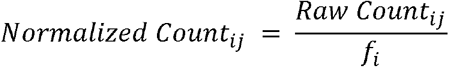

This procedure ensures that the normalized data remains on the scale of the original raw counts, preserving the relationship between the mean and variance across the time-course, which is critical for the ORPHEUS system of equations.

## Downstream Statistical Analyses

To assess differences in phase dispersion between biological groups, we employed two distinct statistical approaches depending on the nature of the comparison.

### Permutation Testing for Homogeneous Populations

When comparing groups where the underlying rhythmic architecture (e.g., cellular amplitude) is expected to be similar, such as distinct hepatocyte subtypes or cells stratified by pathway activity within a single condition, we utilized a permutation-based approach. This method provides a rigorous empirical null distribution.

- **Procedure:** We randomly shuffled the group labels (e.g., cell type or pathway bin) of the individual cells (liver) or subjects (human) while maintaining the original sample sizes.
- **Estimation:** For each iteration, we re-calculated the pseudobulk and variance time-series for the randomized groups and applied ORPHEUS to estimate the gene-wise phase dispersion. We then calculated the difference in median *σ*_*ϕ*_ between the randomized groups.
- **Significance:** This process was repeated 1,000 times to generate an empirical null distribution of differences. The p-value was calculated as the proportion of randomized differences that exceeded the observed difference between the real groups.

### Non-Parametric Testing for Heterogeneous Conditions

When comparing broad biological conditions expected to differ significantly in their fundamental rhythmic properties (e.g., AD vs. CTL), a permutation approach is inappropriate. Shuffling subjects between healthy and diseased states creates artificial populations with high heterogeneity in cellular amplitude (*A*_*c*_). This violates the core assumption of the ORPHEUS model that *A*_*c*_ is effectively constant across the population being analyzed, potentially introducing artifacts into the phase dispersion estimates of the null distribution.

- **Procedure:** Consequently, for the comparison of the AD and CTL excitatory neuron population, we treated the gene-wise phase dispersion estimates as independent observations and compared their distributions directly using a two-sided Wilcoxon rank-sum test. This approach tests for a shift in the central tendency of synchrony across the transcriptome without requiring the model to fit a biologically incoherent mixture of subjects.

### Synchrony-Ranked Gene Set Enrichment Analysis

To test if specific biological pathways were enriched for relatively synchronous or desynchronous genes, we performed a synchrony-ranked gene set enrichment analysis: Genes were ranked based on their ORPHEUS-derived phase dispersion estimates. This ranked list was then used as input for Gene Set Enrichment Analysis (GSEA).

### Pathway-Activity Stratified Synchrony Analysis

To evaluate how cellular physiological states are associated with circadian synchrony, we developed a stratification framework using Gene Set Variation Analysis (GSVA). While the conceptual goal remained consistent, the implementation was tailored to the specific temporal resolution of each dataset.

### Subject-Level Stratification (ROSMAP AD Dataset)

Given the large sample size (n=409) and continuous temporal phase estimates in the human brain data, as well as the fact that we are interested in pathway differences between *subjects*, we performed stratification at the subject level. This was done separately in CTL and AD subjects.

- **Feature Selection:** To ensure pathway scores were not confounded by the subject’s circadian phase, we excluded rhythmic genes, defined as those with a Benjamini-Hochberg adjusted q-value ≥ 0.1 or an amplitude ratio < 0.1, prior to scoring.
- **Scoring and Binning:** We used GSVA to assign pathway scores to each subject, restricting our analysis to pathways containing at least 30 genes (argument “mn.size = 30”). Subjects were then partitioned into tertiles (low, medium, high) for each pathway score.
- **Phase Dispersion Estimation and Empirical Significance** For each pathway, ORPHEUS was performed independently on the “low” and “high” pathway-activity bins. Specifically, rhythmic genes were identified by cosinor regression using all samples in a condition (all samples across pathway-activity bin). Rhythm parameters for these genes were then calculated within each pathway-activity bin and supplied to ORPHEUS to estimate gene-wise phase dispersion. We then assessed the significance of phase dispersion differences between pathway-activity bins: we generated an empirical null distribution by randomly binning subjects into tertiles 1,000 times, recalculating phase dispersion between the “low” and “high” bins for each iteration.

### Cell-Level Stratification (Mouse Liver Dataset)

In the liver dataset, we were limited to a single sample per discrete timepoint and therefore could not stratify samples by pathway score. Instead, we adopted a cell-level stratification strategy:

- **Scoring and Binning:** GSVA scores were assigned to individual cells. To mitigate the inherent noise of single-cell expression, we restricted analysis to pathways with at least 30 genes (argument “mn.size = 30”).
- **Temporal Balancing:** To ensure that stratification did not skew the temporal distribution of the data, cells were binned within each timepoint. For instance, the “top quartile” for a pathway was composed of the top 25% of cells from each of the six timepoints independently. Because stratification occurred within timepoints, rhythmic genes were retained.
- **Phase Dispersion Estimation and Empirical Significance:** For each pathway, ORPHEUS was performed independently on the “low” and “high” pathway-activity bins. Specifically, we estimated phase dispersion for the rhythmic genes identified by BooteJTK. Rhythm parameters for these genes were calculated within each pathway-activity bin and supplied to ORPHEUS to estimate gene-wise phase dispersion. We then assessed the significance of phase dispersion differences between pathway-activity bins: we generated an empirical null distribution by randomly binning subjects into quartiles 1,000 times, recalculating phase dispersion between the “low” and “high” bins for each iteration.

## Data and Code Availability

The ORPHEUS R package and custom scripts are available at GitHub: https://github.com/ranafi/Orpheus. The human brain snRNA-seq data from the ROSMAP cohort are available via the Synapse repository https://www.synapse.org/#!Synapse:syn3219045.

Access requires a data use certificate: https://adknowledgeportal.synapse.org/Data%20Access. The mouse SCN time-course and the mouse liver time-course data were obtained from the Gene Expression Omnibus with identifiers GSE117295, and GSE292219, respectively.

## Acknowledgements

This work was supported by NIA R01AG068577 (R.C.A.). ROSMAP was supported by NIA P30AG10161, NIH P30AG72975, NIA R01AG15819, NIA R01AG17917, NIH U01AG46152, and NIH U01AG61356. We used data from the AD Knowledge Portal (www.synapse.org).

